# VPAC1 couples with TRPV4 channel to promote calcium-dependent gastric cancer progression

**DOI:** 10.1101/360404

**Authors:** Bo Tang, Jilin Wu, Michael X. Zhu, Xuemei Sun, Jingjing Liu, Rui Xie, Tobias Xiao Dong, Yufeng Xiao, John M. Carethers, Shiming Yang, Hui Dong

## Abstract

Although VPAC1 and its ligand vasoactive intestinal peptide (VIP) are important in gastrointestinal physiology, their involvements in progression of gastrointestinal tumor have not been explored. Here, we found that higher expression of VIP/VPAC1 was observed in gastric cancer compared to the adjacent normal tissues. The increased expression of VIP/VPAC1 in gastric cancer correlated positively with invasion, tumor stage, lymph node, distant metastases, and poor survival. Moreover, high expression of VIP and VPAC1, advanced tumor stage and distant metastasis were independent prognostic factors. VPAC1 activation by VIP markedly induced TRPV4-mediated Ca^2+^ entry, and eventually promoted gastric cancer progression in a Ca^2+^ signaling-dependent manner. Inhibition of VPAC1 and its signaling pathway could block the progressive responses. VPAC1/TRPV4/Ca^2+^ signaling in turn enhanced the expression and secretion of VIP in gastric cancer cells, enforcing a positive feedback regulation mechanism. Taken together, our study demonstrate that VPAC1 is significantly overexpressed in gastric cancer and VPAC1/TRPV4/Ca^2+^ signaling axis could enforce a positive feedback regulation in gastric cancer progression. VIP/VPAC1 may serve as potential prognostic markers and therapeutic targets for gastric cancer.

## Introduction

Gastric cancer (GC) is the second leading cause of cancer-related death worldwide (Siegel RL *et al*, 2016). Despite important advances in the understanding of pathology and molecular mechanisms, the prognosis of GC remains poor due to the early invasion and metastasis (Tan P & Yeoh KG, 2015; Badgwell B *et al*, 2016). Currently, there are no specific molecular signatures of gastric cancer metastasis and progression. Thus, it is urgent to explore novel diagnostic markers and mechanisms responsible for the development and progression of GC.

VPAC1, a member of the G-protein-coupled receptor (GPCR) superfamily, is mainly activated by vasoactive intestinal peptide (VIP), a ligand of VPAC1 that belongs to the pituitary adenylyl cyclase-activating polypeptide family (Vacas E *et al*, 2013; Vacas E *et al*, 2013). Aberrant expression of VPAC1 is implicated in several human diseases, such as autoimmune disease, neurological disease and carcinogenesis (Seoane IV *et al*, 2016; White CM *et al*, 2010; Valdehita A *et al*, 2009; (Valdehita A *et al*, 2007). The role of VPAC1 and VIP in tumor progression has been gradually recognized, and they are frequently overexpressed in a variety of malignancies (Reubi JC et al, 2000; Fernández-Martínez AB et al, 2012; Liu S et al, 2014). However, the role of VPAC1 in gastric cancer progression remains unknown.

VPAC1 activation induces not only an increase in intracellular cAMP in many types of cells, but also a rise in cytoplasmic free Ca^2+^ ([Ca^2+^]_cyt_) (Dickson L *et al*, 2009; Zhang S *et al*, 2010; Herrera JL *et al*, 2009). [Ca^2+^]_cyt_ elevation is a mechanism well suited to the rapid transmission of signals from tumor microenvironment into cellular responses (Berridge MJ *et al*, 2003; Monteith GR *et al*, 2012), and an ubiquitous mechanism that controls cell survival, proliferation, motility, differentiation, and cancer progression (Fiorio Pla A *et al*, 2012; Wei C *et al*, 2009).

Differences in Ca^2+^ signature, which are key to specific Ca^2+^-dependent cellular responses, often rely upon complex spatiotemporal variations in [Ca^2+^]_cyt_ (Berridge MJ *et al*, 2003; Evans AM *et al*, 2016). A major determinant of these is the participation of distinct Ca^2+^-permeable channels (Hofer AM & Brown EM, 2003; Déliot N & Constantin B, 2015), among which transient receptor potential vanilloid 4 (TRPV4), a member of the transient receptor potential (TRP) superfamily, is able to regulate a variety of Ca^2+^-dependent cellular functions in gastrointestinal tract (Holzer P, 2011; D’Aldebert E *et al*, 2011). However, except for a few studies to address its involvement in inflammatory bowel diseases (Holzer P, 2011; D’Aldebert E *et al*, 2011), nothing is currently known about the pathological role of TRPV4 in gastrointestinal cancers.

In the present study, we demonstrate for the first time an overexpression of VPAC1 in human gastric cancer, which is closely correlated with advanced clinicopathological features and poor survival of gastric cancer patients. We also reveal that VPAC1 activation promotes migration and invasion of GC cells through TRPV4 channel-dependent Ca^2+^ entry, which in turn augments VIP expression and secretion to drive several downstream signaling cascades to promote GC progression. Enhanced expressions of VPAC1 have potentially diagnostic and prognostic significance for gastric cancer, and targeted inhibition of VPAC1 expression and its downstream signaling could attenuate metastasis of gastric cancer, which may be a novel candidate for gastric cancer diagnosis and therapy.

## Results

### Enhanced expression of VIP and VPAC1 in primary gastric cancer correlates with metastasis and poor prognosis

We first measured the expression level of VPAC1 in gastric cancer cell lines, normal gastric epithelial cell GES-1 and gastric cancer tissues. As shown in Appendix Fig S1A and B, both mRNA and protein expression levels of VPAC1 were much higher in gastric cancer cells than in normal gastric GES-1 cells. Consistently, VPAC1 was significantly upregulated in gastric cancer tissues compared with their matched adjacent non-tumor tissues (Appendix Fig S1C and D). We further examined the expression of VIP and VPAC1 in biopsy samples from 238 GC patients using immunohistochemistry (Fig 1A). Semi-quantitative analysis showed an obvious upregulation of VIP and VPAC1 expression in GC tissues compared with adjacent non-tumor tissues (Fig 1B and C). Furthermore, VIP and VPAC1 were expressed at higher levels in the metastasis-positive group than those in the non-metastasis group (*P*<0.001) and were related to advanced TNM stage (*P*<0.0001) (Fig 1D and E). We further evaluated the correlation between VIP/VAPC1 levels and patient clinicopathological characteristics, and found that high expression of VIP and VPAC1 was significantly associated with deeper tumor invasion, lymph node metastasis, advanced tumor stage and distant metastasis (Appendix Table S1). Additionally, Kaplan-Meier survival analysis showed that the high expression of VIP and VPAC1 strongly correlated with a decreased survival time (*P*<0.0001) (Fig 1F and G), consistent with the results extracted from KM Plotter database (Appendix Fig S2). Notably, short survival was observed in the group with high expression of both VIP and VPAC1, but long survival in the group with low expression of both proteins (Fig 1H). To further explore the role of VPAC1 and VIP in predicting gastric cancer prognosis, we performed univariate and multivariate analyses of clinical follow-up data. We found that deeper tumor invasion, lymph node metastasis, advanced tumor stage, distant metastasis and higher expression of VIP and VPAC1 were significantly associated with poorer survival on univariable analysis (Appendix Table S2). Multivariable Cox regression analysis indicated that high VIP (HR=1.76; 95%CI, 1.16 to 2.68; *P*=0.008) and VPAC1 expression (HR=2.09; 95%CI, 1.33 to 3.31; *P*=0.002), advanced tumor stage (HR=2.47; 95%CI, 1.49 to 4.10; *P*<0.001) and distant metastasis (HR=1.61; 95%CI, 1.35 to 2.89; *P*=0.003) were independent prognostic factors (Appendix Table S2). Taken together, these data indicated that VIP/VPAC1 significantly contribute to gastric cancer metastasis and progression, implicating them as potential prognostic biomarker in gastric cancer.

**Fig. 1.**
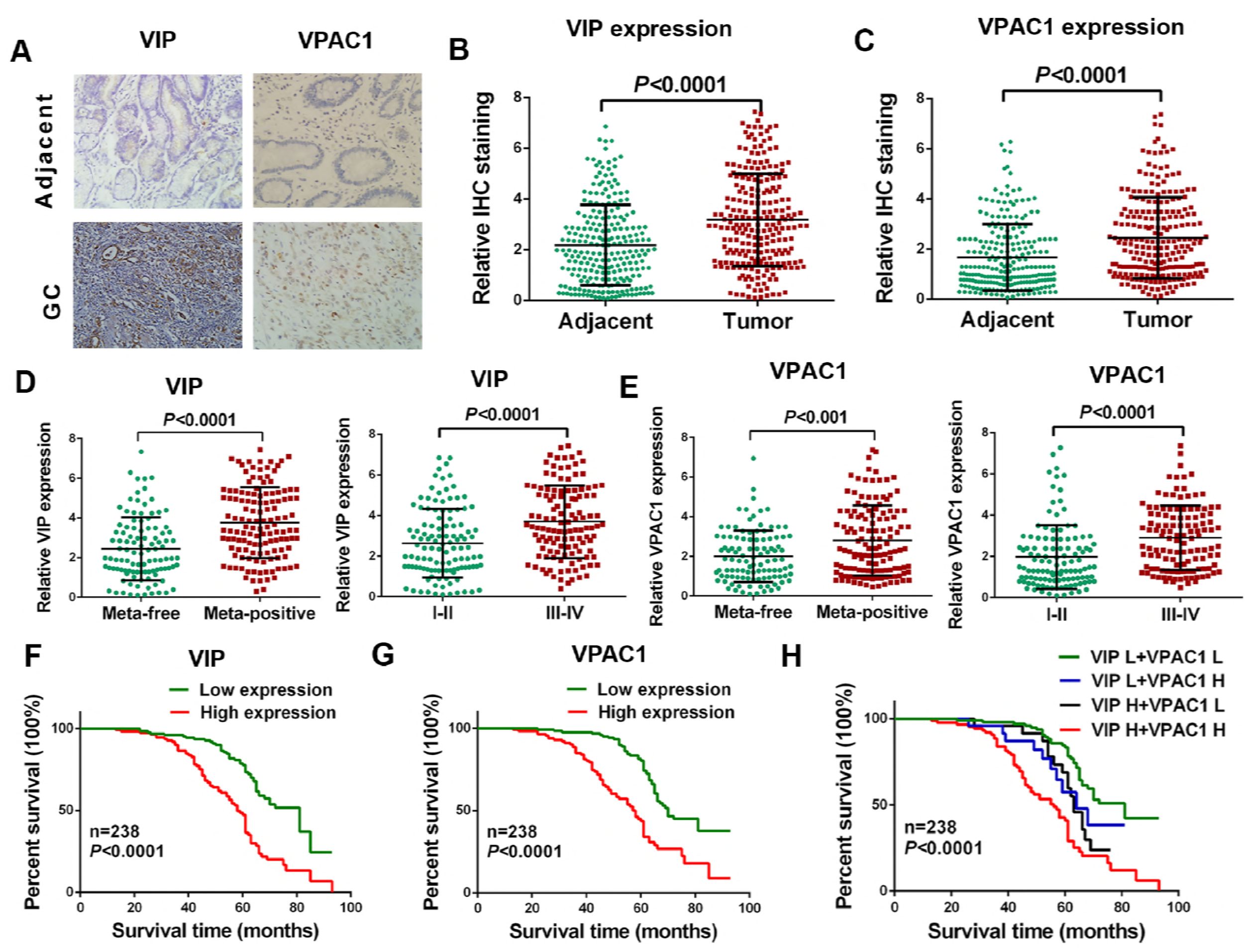
Upregulation of VIP and VPAC1 correlates with high metastatic potential and poor prognosis of human GC. (A) Immunohistochemical staining for VIP and VPAC1 expression in human GC tissues and matched adjacent normal tissues. (B, C) VIP and VPAC1 expression levels were scored with semiquantitative immunohistochemical (IHC) analysis. (D) The relation between VIP expression and lymphatic metastasis/TNM stage in GC patients. (E) The relation between VPAC1 expression and lymphatic metastasis/TNM stage in GC patients. (F) Kaplan-Meier curves comparing overall survival in GC patients with low and high levels of VIP expression (n=238; *P*<0.0001, log-rank test). (G) Kaplan-Meier curves comparing overall survival in GC patients with low and high levels of VPAC1 expression (n=238; *P*<0.0001, log-rank test). (H) Kaplan-Meier analyses of overall survival according to the combination of the above two indices (n=238; *P*<0.0001).

### VIP enhances migration and invasion of gastric cancer cells *in vitro* and metastasis *in vivo* through activation of VPAC1

We found that VIP enhanced the migration and invasion of GC cells, which was attenuated by knockdown of VPAC1 (Fig 2A and B and Appendix Fig S3A and B). Furthermore, VIP significantly increased the lung metastasis and peritoneal dissemination, while VPAC1 knockdown markedly attenuated the effect of VIP (Fig 2C and D). Our immunohistochemistry and western blot analysis confirmed the effective knockdown of VPAC1 in the tumor nodules (Appendix Fig S4A and B). Taken together, these results strongly suggest that VIP promotes the migration, invasion and metastasis of GC through VPAC1.

**Fig. 2.**
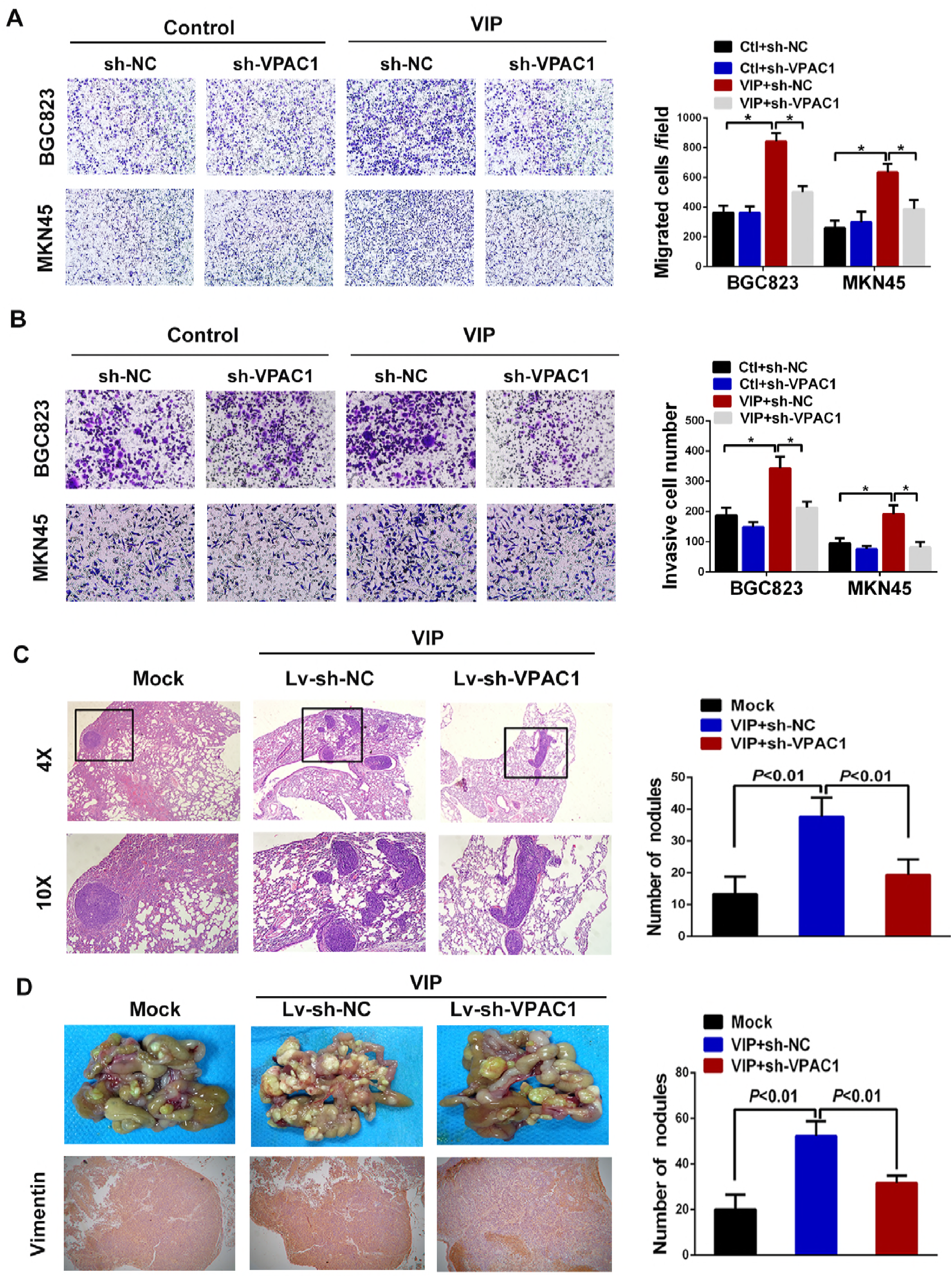
VIP enhances gastric cancer cells invasion and metastasis through VPAC1. (A) Boyden chamber migration assay. Cells treated as indicated were allowed to migrate for 24 h across the membrane in Boyden chambers. Cells that migrated across the filter were counted in 20 fields per well. Each experiment was performed at least three times, \**P*<0.05. (B) Representative micrographs of the transwell matrix penetration assay showing the invasiveness of VPAC1-silenced cells compared with vector controls stimulated by VIP. Data shown in histogram are summary of three independent experiments, \**P*<0.05. (C) A pulmonary metastasis model was established in nude mice by injecting into the tail vein with MKN45 cells. VIP or PBS was injected through tail vein once a day. After 12 weeks, the lungs were isolated for hematoxylin and eosin staining. The black squares in the sections show the tumor foci (*left panel*), and histogram shows the calculated number of metastasis nodules in pulmonary tissues (*right panel*). (D) A peritoneal dissemination model was established in nude mice by inoculating the abdominal cavity with MKN45 cells and VIP. Representative micrographs (*left panel*) and the calculated number of tumor nodules (*right panel*) are shown.

### VPAC1 activation induces Ca^2+^ signaling to promote migration and invasion of gastric cancer cells

VPAC1 is also coupled to phospholipase Cβ (PLCβ) and Ca^2+^ signaling that is critical for the migration and invasion of cancer cells (Chen YF *et al*, 2013, 2016). Indeed, VIP induced [Ca^2+^]_cyt_ increases in MKN45 cells in a concentration-dependent manner (Fig 3A), which was abolished by VPAC1 knockdown (Fig 3B and Appendix Fig S5A and B). We further confirmed that VPAC1 mediated the VIP-induced Ca^2+^ signal by stably expressing the receptor in HEK293 cells since VIP induced a [Ca^2+^]_cyt_ rise only in VPAC1-expressing HEK293 cells (Fig 3C). Moreover, U73122, a PLC inhibitor, significantly attenuated the VIP-induced [Ca^2+^]_cyt_ rise (Fig 3D). Finally, BAPTA-AM, a [Ca^2+^]_cyt_ chelator, significantly attenuated VIP-induced migration and invasion of GC cells (Appendix Fig S6A-F). Collectively, our data demonstrate that VIP activation of VPAC1 induces Ca^2+^ signaling, which can play a critical role in the progression of GC cells.

**Fig. 3.**
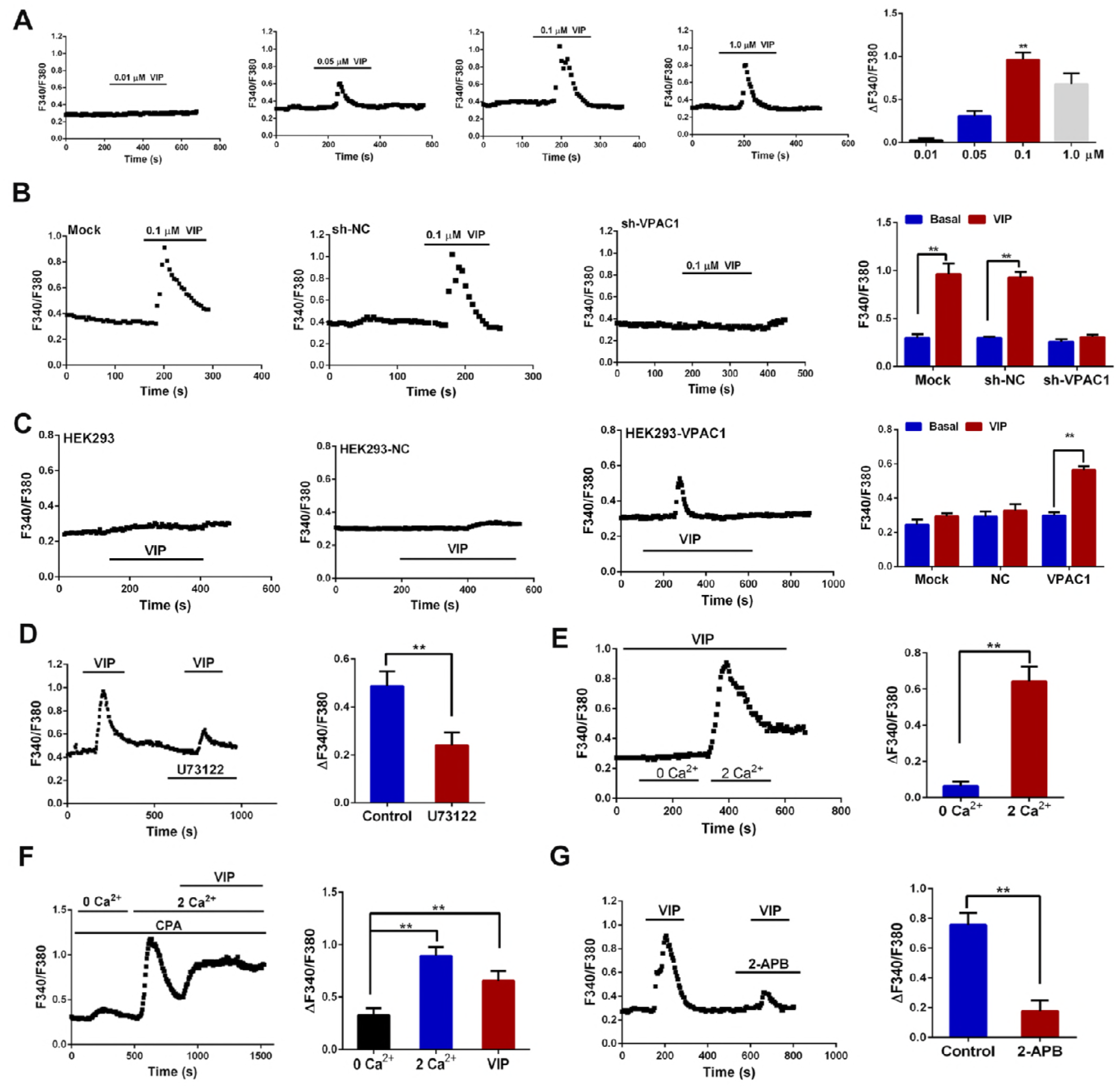
VIP induces an increase in cytosolic Ca^2+^ *via* extracellular Ca^2+^ entry through activation of VPAC1. (A) Time courses of Fura-2 ratio ([Ca^2+^]_cyt_) changes in MKN45 cells induced by different concentrations of VIP. Histogram at the right shows the summary data for peak changes of Fura-2 fluorescence ratio (the difference between the basal and peak values) induced by VIP, \*\**P*<0.01, compared to 0.1 µM VIP. (B) Effect of VPAC1 knockdown on VIP-induced [Ca^2+^]_cyt_ in MKN45 cells. (C) Effects of VIP on HEK293 cells that expressed or not VPAC1. HEK293 cells were mock transfected or transfected with a control vector (NC) or VPAC1 and stimulated with 0.1 µM VIP. (D) Time courses of [Ca^2+^]_cyt_ changes in MKN45 cells induced by VIP (0.1 µM) in the absence and presence of U73122 (10 μM). (E) Time course of [Ca^2+^]_cyt_ changes in MKN45 cells induced by 0.1 μM VIP in Ca^2+^-free and then normal Ca^2+^ PSS. (F) In the presence of 10 μM CPA, a change of extracellular Ca^2+^ concentration from 0 to 2 mM induced a rise in [Ca^2+^]_cyt_,.reflective of store-operated Ca^2+^ entry (SOCE). Further addition of 0.1 μM VIP evoked an additional substantial increase in [Ca^2+^]_cyt_ despite store depletion and active SOCE. (G) Time course of [Ca^2+^]_cyt_ changes in MKN45 cells induced by VIP in the absence and presence of 2-APB (100 μM). Summary data in histograms represent mean ± SEM of 40-50 cells for each group. \*\**P*<208:648 0.01.

### VPAC1 activation induces Ca^2+^ signaling via extracellular Ca^2+^ entry rather than internal Ca^2+^ release in gastric cancer cells

Since [Ca^2+^]_cyt_ rise may result from internal Ca^2+^ release and/or extracellular Ca^2+^ entry (Zhu MX *et al*, 2016), we further determined the Ca^2+^ source of VIP-evoked [Ca^2+^]_cyt_ rise in GC cells. When VIP was first applied in a Ca^2+^-free solution, it did not elicit any responses; however, reintroduction of Ca^2+^ induced a marked [Ca^2+^]_cyt_ increase (Fig 3E and Appendix Fig S5C). Moreover, in the presence of CPA, a specific inhibitor of sarcoendoplasmic reticulum Ca^2+^-ATPase (SERCA), VIP evoked an additional substantial increase in [Ca^2+^]_cyt_ on the top of CPA-induced store-operated Ca^2+^ entry (SOCE) (Fig 3F), indicating that VIP induced Ca^2+^ entry via a mechanism that is different from SOCE. Nonetheless, the VIP-induced [Ca^2+^]_cyt_ rise was attenuated by the pretreatment with 2-aminoethoxydiphenyl borate (2-APB, 100 µM) (Fig 3G and Appendix Fig S5D), a nonspecific inhibitor of TRP channels, suggesting their possible involvements in VIP-induced Ca^2+^ entry.

### VPAC1 activation induces Ca^2+^ entry through TRPV4 channels in gastric cancer cells

We then screened a panel of Ca^2+^-permeable channels in GC cells, and found that SKF96365 (10 µM), a commonly used blocker for TRPC channels, did not alter VIP-induced [Ca^2+^]_cyt_ rise (Fig 4A). L-type voltage-gated Ca^2+^ channel blocker, nifedipine (10 µM), did not affect VIP-induced [Ca^2+^]_cyt_ (Fig 4A) and neither did the Na^+^/Ca^2+^ exchanger 1 inhibitor, SN-6 (Fig 4A). However, the VIP-induced [Ca^2+^]_cyt_ rise was abolished by a selective TRPV4 antagonist, RN1734 (10 µM) (Fig 4A, Appendix Fig S7A), suggesting that TRPV4 is a likely molecular candidate that mediates VIP/VPAC1-evoked Ca^2+^ entry.

**Fig. 4.**
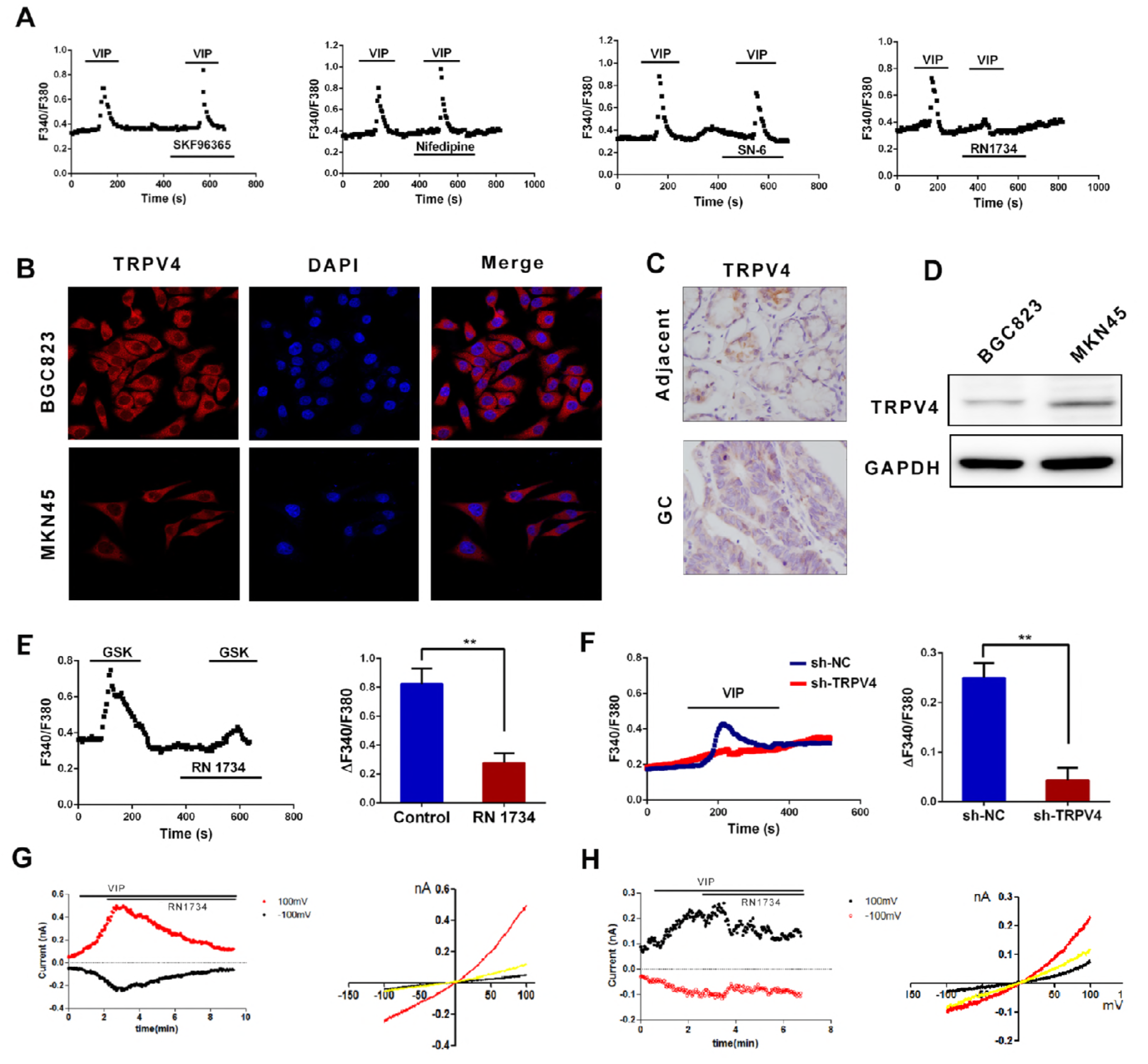
VIP induces Ca^2+^ entry through TRPV4. (A) Effects of SKF96365 (10 μM), nifedipine (10 μM), SN-6 (10 μM), and RN1734 (10 μM) on VIP-induced [Ca^2+^]_cyt_ changes in MKN45 cells. (B) Representative images of immunofluorescence staining of TRPV4 proteins in GC cells. (C) Representative photographs showing expression of TRPV4 proteins in GC and adjacent normal tissues in human biopsy samples using immunohistochemical analysis. (D) Western blot analysis of TRPV4 protein expression in BGC823 and MKN45 cells. (E) Effect of RN1734 (10 μM) on GSK1016790A-induced [Ca^2+^]_cyt_ in MKN45 cells (\*\**P*<0.01, n=30). (F) Effect of TRPV4 knockdown on VIP-induced [Ca^2+^]_cyt_ rise in MKN45 cells (\*\**P*<0.01, n=40). (G, H) Representative traces of whole-cell currents evoked by VIP (0.1 µM) and their suppression by RN1734 (10 µM) in MKN45 GC cells (G) and normal gastric GES-1 cells (H). Left panels show time courses of currents at +100 and −100 mV. Right panels show current-voltage relationships obtained using voltage ramps under basal (*black line*), at the peak of the response to VIP (*red line*) and after the suppression by RN1734 (*yellow line*).

By immunofluorescence staining, we found TRPV4 expression in GC cells (Fig 4B). We also confirmed the expression of TRPV4 in human normal gastric tissues, GC tissues and cells (Fig 4C and D). A selective TRPV4 agonist, GSK1016790A (1 µM), did not alter [Ca^2+^]_cyt_ in the Ca^2+^-free solution, but induced a marked [Ca^2+^]_cyt_ rise in Ca^2+^-containing solutions (Appendix Fig S7B and C), which was strongly attenuated by RN1734 (Fig 4E and Appendix Fig S7D) and the knockdown of TRPV4 expression (Fig 4F). VIP also evoked outwardly rectifying currents in MKN45 cells (Fig 4G), and to a lesser extent in GES-1 cells (Fig 4H), which were both inhibited by RN1734, confirming that VAPC1 activation by VIP induces Ca^2+^ entry through TRPV4 channels.

### VPAC1 activation induces Ca^2+^ entry via TRPV4 channels in a DAG-dependent manner

To elucidate the detail mechanisms by which VIP/VPAC1 activates TRPV4 channels, we first tested if VIP/VPAC1 directly interacts with TRPV4 in GC cells. However, VIP did not affect TRPV4 expression (Appendix Fig S8A and B), and no interaction was observed between VPAC1 and TRPV4 channels (Appendix Fig S8C). Since VPAC1 is coupled to PLCβ stimulation which produces diacylglycerol (DAG) that can activate TRPC channels (Leuner K *et al*, 2010; Storch U *et al*, 2017), we examined whether 1-oleoyl-2-acetyl-sn-glycerol (OAG), a DAG analog, could induce Ca^2+^ responses in GC cells. Indeed, OAG elevated [Ca^2+^]_cyt_ in GSK-responsive GC cells (Fig 5A and D), which was abolished by RN1734 and TRPV4 knockdown (Fig 5B and E). The OAG-induced [Ca^2+^]_cyt_ rise was dependent on extracellular Ca^2+^ entry (Fig 5C and D). Moreover, in TRPV4-expressing HEK293 cells but not parental HEK293 cells, OAG induced [Ca^2+^]_cyt_ increase, which was blocked by RN1734 (Fig 5F). In GC cells, the VIP-induced [Ca^2+^]_cyt_ rise was strongly reduced by the expression of DAG kinase β (DGKβ), an enzyme that decreases the intracellular DAG level by converting DAG to phosphatidic acid (Fig 5G), but augmented by the treatment of RHC80267, a DAG lipase inhibitor that causes accumulation of endogenous DAG (Fig 5H). Taken together, these results clearly indicate that VIP/VPAC1 activates TRPV4 channels via PLC generation of DAG.

**Fig. 5.**
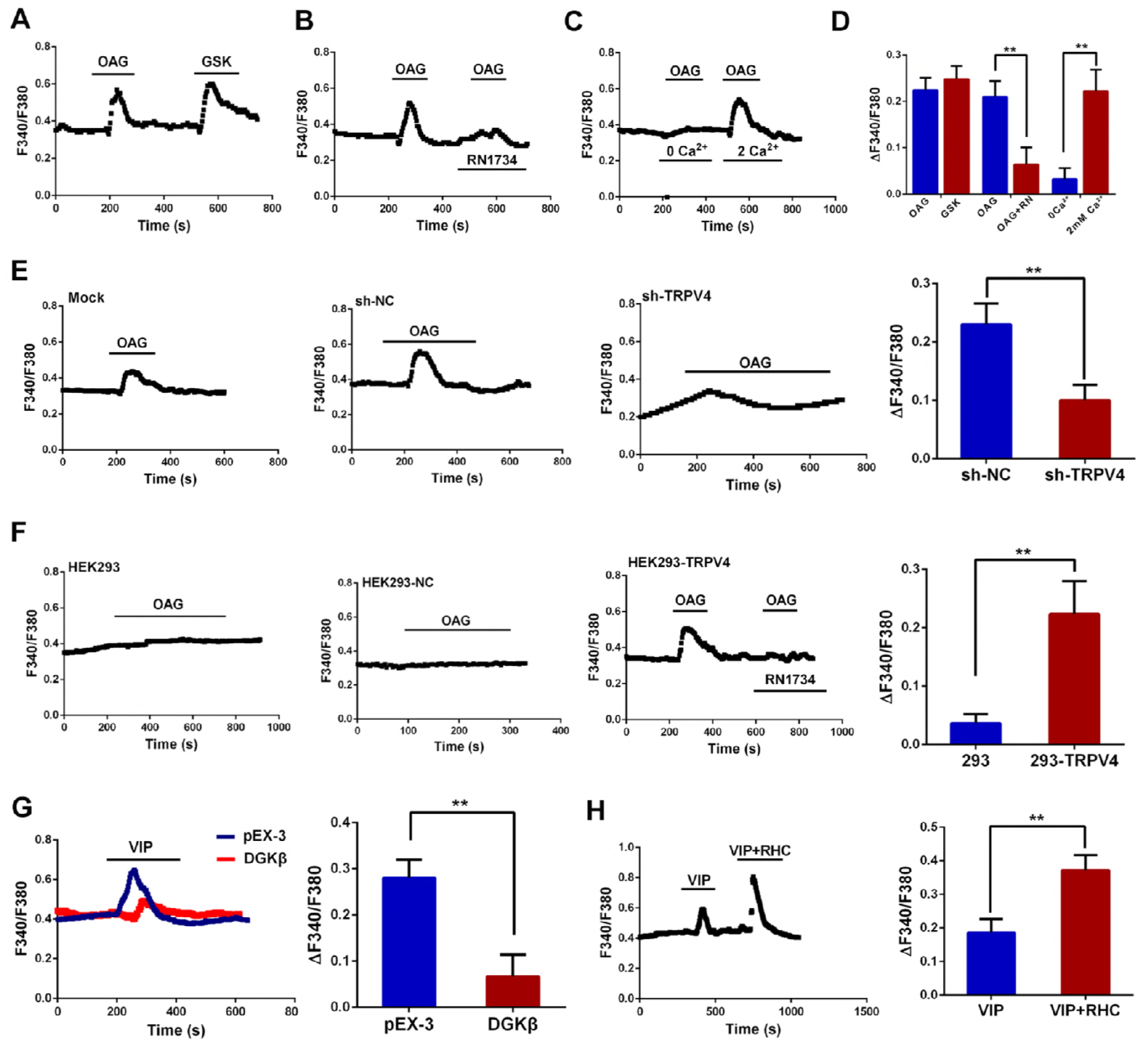
VIP activates TRPV4-mediated Ca^2+^ entry in a DAG-dependent manner. (A) Representative trace of Fura-2 ratio ([Ca^2+^]_cyt_) changes induced by 100 μM OAG in GSK-responsive TRPV4-expressing cells. (B, C) The OAG-evoked Ca^2+^ transients were blocked by the pretreatment of 10 μM RN1734 (B) and Ca^2+^-free extracellular solution (C). (D) Summary data of 40-50 cells for each group. \*\**P*<0.01. (E) The knockdown of TRPV4 expression attenuated [Ca^2+^]_cyt_ rise evoked by OAG. (F) OAG (100 µM) induced RN1734-sensitive Ca^2+^ transients in HEK293 cells that expressed TRPV4, but not the untransfected or vector-transfected HEK293 cells. (G) Effects of transfection of DGKβ on VIP-induced Ca^2+^ transients in GC cells.(H) Effects of RHC80267 treatment on VIP-induced Ca^2+^ transients in GC cells. For (E)-(H), summary data include 40-50 cells for each group. \*\**P*<0.01.

### VIP/VPAC1 promotes gastric cancer progression through TRPV4 -mediated Ca^2+^ signaling pathways

RN1734 markedly decreased VIP-induced migration (Appendix Fig S9A and B) and invasion of GC cells (Appendix Fig S9C). Furthermore, TRPV4 knockdown significantly decreased VIP-induced migration and invasion (Fig 6A). Similarly, TRPV4 knockdown greatly reduced VIP-induced lung metastases (Fig 6B) and peritoneal dissemination in mice (Fig 6C). Together, these results suggest that TRPV4 plays a critical role in VIP/VPAC1 promotion of GC invasion and metastases.

**Fig. 6.**
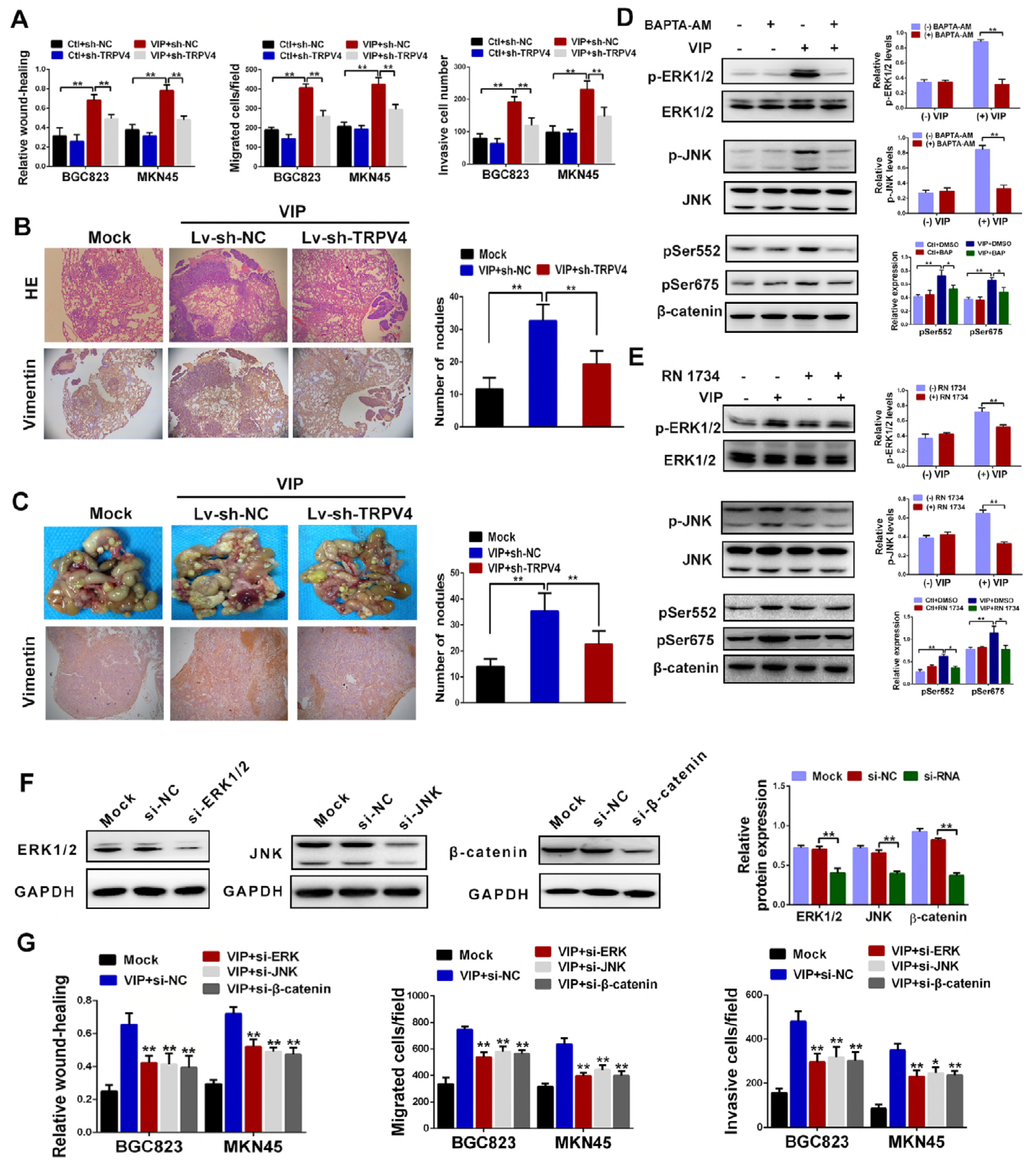
VIP stimulation of GC progression is dependent on TRPV4-mediated Ca^2+^ signaling. (A) Histograms of summary results of wound healing, migration and transwell matrix penetration assays showing the effects of TRPV4 knockdown on VIP-induced migration and invasion of GC (BGC823 and MKN45) cells. Data are representative of three independent experiments, \*\**P*<0.01. (B, C) In pulmonary metastasis (B) and peritoneal dissemination (C) models using MKN45 cells to inoculate nude mice, TRPV4 knockdown reduced the number of tumor nodules. Representative micrographs (*left panels*) and summary for the number of tumor nodules (*right panels*) are shown. \*\**P*<0.01. (D) *Left panels*: representative immunoblots showing the effects of VIP on ERK1/2, JNK, and β-caternin (pSer552 and pSer675) activation in MKN45 cells and their suppression by BAPTA-AM. *Right panels*: summary data from semiquantitative analyses from three independent experiments, \**P*<0.05, \*\**P*<0.01. (E) TRPV4 inhibitor RN1734 (10 µM) attenuated VIP-induced phosphorylation of ERK1/2, JNK, and βcaternin (Ser552 and Ser675) in MKN45 cells. Representative immunoblots (*left panels*) and summary for three independent experiments (*right panels*) are shown. \**P*<0.05, \*\**P*<0.01. (F) Knockdown efficiencies of ERK1/2, JNK, and β-caternin in MKN45 cells by the corresponding siRNA’s, determined by western blot analysis. All experiments were performed at least three times, \*\**P*<0.01. (G) The siRNA knockdown of ERK1/2, JNK, or β-caternin expression in BGC823 and MKN45 cells reduced VIP-induced migration and invasion in wound healing, migration and transwell matrix penetration assays.

To identify the downstream signaling of TRPV4 activation induced by VPAC1 activation, we evaluated Ca^2+^-dependent activation of β-catenin, ERK1/2, and JNK, which have been suggested to be important for GC progression (Xiao G *et al*, 2015; Song W *et al*, 2014; Ma Y *et al*, 2016).

We showed that VIP treatment increased phosphorylation of β-catenin, ERK1/2 and JNK in a time-dependent manner (Appendix Fig S10A), which was significantly attenuated by VPAC1 knockdown (Appendix Fig S10B), BAPTA-AM (Fig 6D) or RN1734 (Fig 6E). To examine the roles of these signaling pathways in VIP-promoted GC progression, we knocked down ERK1/2, JNK and β-catenin, which parallelly led to significant decreases in both protein expression and VIP-induced migration and invasion of GC cells (Fig 6F and G). Therefore, TRPV4/Ca^2+^-mediated β-catenin, ERK1/2, and JNK pathways play key roles in VIP/VPAC1-promoted GC progression.

### VPAC1/TRPV4/Ca^2+^ signaling enhances VIP expression and secretion in gastric cancer cells

Since serum VIP concentration was increased in patients with gastrointestinal cancers (Hejna M *et al*, 2001), we examined whether GC cells could secrete VIP to produce an autocrine regulation during GC progression. Interestingly, we found that VIP could induce VIP mRNA expression in a time-dependent manner (Fig 7A), which was attenuated by the knockdown of VPAC1 (Fig 7B) or TRPV4 (Fig 7C), or BAPTA-AM (Fig 7D), indicating that VIP induces VIP mRNA expression in GC cells through VPAC1/TRPV4/Ca^2+^ signaling pathway. We also found that VIP protein secretion in culture medium was significantly inhibited by the knockdown of TRPV4, or by BAPTA-AM (Fig 7E). To further elucidate the underlying mechanisms, we tested cAMP-responsive element binding protein (CREB) and nuclear factor of activated T-cells (NFAT), two transcription factors known to be essential for Ca^2+^-mediated gene regulation (Yin Y *et al*, 2016; Hogan PG, 2017). We found that the knockdown of CREB but not NFAT attenuated the VIP-induced increase in VIP expression (Fig 7F). Similarly, serum VIP secretion was reduced by the knockdown of VPAC1 or TRPV4 in GC xenografts *in vivo* (Fig 7G), and VIP expression in the tumor tissues of GC xenografts was also significantly decreased by the knockdown of VPAC1 or TRPV4 (Fig 7H and I). Furthermore, the expression levels of VIP were positively correlated with those of VPAC1 in GC specimens (Fig 7J), and serum VIP concentrations in GC patients were higher than those in normal individuals (Fig 7K), which is consistent with previous studies (Hejna M et al, 2001). VIP knockdown significantly inhibited GC invasion and metastasis, which were further reduced by knockdown of both VIP and VPAC1 (Appendix Fig S11). More importantly, survival was improved in mice after VIP or VPAC1 knockdown (Appendix Fig S12), which is consistent with the survival results of GC patients. Taken together, these results reveal a positive feedback mechanism of VIP/VPAC1/TRPV4/Ca^2+^ signaling via CREB to promote GC progression by enhancing VIP expression and secretion.

**Fig. 7.**
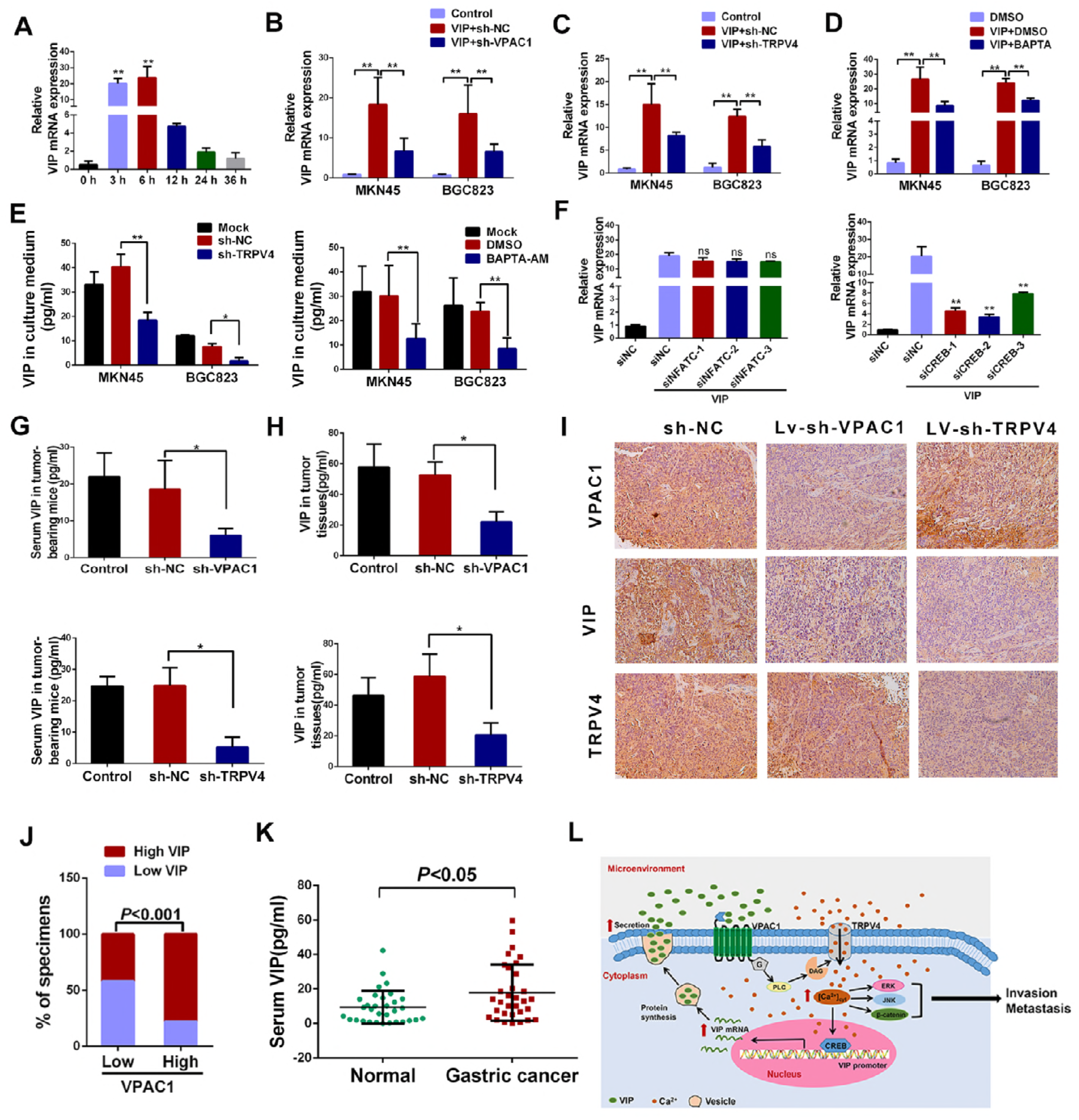
VPAC1/TRPV4-mediated Ca^2+^ signaling regulates VIP expression and secretion. (A) VIP mRNA expression was increased in response to VIP (0.1 µM) within 3-6 h and then gradually declined. Following the addition of VIP, MKN45 cells were collected at times indicated (0, 3, 6, 12, 24, and 36 h) and total RNA was prepared and subjected to RT and qPCR. Data represent mean ± SD from at least three independent experiments. \*\**P*<0.01 vs. control at 0 h. (B-D) MKN45 and BGC823 cells were transfected with sh-VPAC1 (B) or sh-TRPV4 (C), or treated with BAPTA-AM (D). The effect of VIP on VIP mRNA expression levels was then determined using RT followed by qPCR. \*\**P*<0.01. (E) VIP levels in culture medium and their suppression by TRPV4 knockdown (*left panel*) and BAPTA-AM treatment (*right panel*). \**P*<0.05, \*\**P*<0.01. (F) In MKN45 cells, siRNA knockdown of CREB (*right panel*), but not NFAT (*left panel*) attenuated the VIP-induced expression of its own mRNA, as determined by RT followed by qPCR. (G) MKN45 cells transfected with LV-sh-VPAC1 (*upper panel*), LV-sh-TRPV4 (*lower panel*), or the control vector were injected into tail veins of nude mice. At 12 weeks after the injection, mouse serum was collected and VIP concentration in the serum was determined using ELISA. \**P*<0.01. (H) The same mice as in (G), but tumor tissues were collected and VIP concentrations in the tissues were determined using ELISA. \**P*<0.01. (I) Immunohistochemical staining for VIP, VPAC1 and TRPV4 expression in tumor grafts from mice in (G) and (H). (J) Percentages of tissue specimens with low or high expression of VIP categorized by those that had low or high expression of VPAC1. (K) Serum VIP concentrations of human GC patients and normal individuals, as determined by ELISA. (L) Schematic representation of VIP/VPAC1/TRPV4/Ca^2+^ signaling pathway in GC progression. VIP binds to VPAC1 receptor, which subsequently activates TRPV4 channel in a DAG-dependent manner to mediate Ca^2+^ entry. The increase in [Ca^2+^]_cyt_ activates several signaling pathways to promote migration and invasion of GC cells. Additionally, Ca^2+^ can also activate transcription factor CREB, which in turn promotes the expression and secretion of VIP, indicating a positive feedback regulation in GC progression.

## Discussion

Tumor metastasis is a complicated process with diverse mechanisms, and it could be promoted by the production of cytokines, chemokines, growth factors, neuropeptides and proteinases (Chen F *et al*, 2015). In the present study, we identified that VIP and VPAC1 were extensively overexpressed in human primary GC tissues and cell lines, which were positively correlated with advanced clinical stage, lymphatic metastasis, and poor survival, suggesting VIP/VPAC1 played critical roles in regulating GC metastasis and progression. Mechanistically, VPAC1 activation promotes migration and invasion of GC cells through TRPV4 channel-dependent Ca^2+^ entry, which in turn augments VIP expression and secretion to form a feedback regulation to worsen GC progression. Importantly, enhanced expressions of VPAC1 have potentially diagnostic and prognostic significance for gastric cancer. Taken together, these findings indicate that VPAC1 serves as an important regulator of GC metastasis and progression, which may be a novel candidate for gastric cancer diagnosis and therapy.

Previous studies showed that Ca^2+^ signaling elicits many biological responses, and aberrant Ca^2+^ signaling is involved in different pathogenesis (Berridge MJ *et al*, 2003; Galione A & Churchill GC, 2002). VIP stimulation and VPAC1 activation are well recognized to trigger Ca^2+^ signaling (Li D *et al*, 2013; Hagen BM *et al*, 2006), but it remains totally unknown whether VIP-mediated Ca^2+^ signaling plays a role in gastrointestinal malignancy. Here we demonstrated for the first time that inhibition of TRPV4-mediated Ca^2+^ entry effectively inhibited VIP/VPAC1-stimulated GC progression. Previous studies showed that GPCR activation could promote DAG/PKC to stimulate activities of TRPV1 and TRPV4 channels (Woo DH *et al*, 2008). Consistently, our results indicated that VIP activation of VPAC1 enhanced TRPV4 channels via PLC-induced DAG generation in GC cells. Since [Ca^2+^]_cyt_ rise is well known to activate ERK1/2, JNK and βcatenin in different cell types (Andrikopoulos P *et al*, 2015; Favia A *et al*, 2014; Thrasivoulou C *et al*, 2013), we found that most of these downstream effecter kinases were also activated by TRPV4-mediated [Ca^2+^]_cyt_ rise in GC cells. Taken together, we identified a novel coupling of VPAC1 and TRPV4 channels and subsequent activation of Ca^2+^-dependent kinases, which may play a major role in GC progression.

It is worthy to note that serum VIP concentrations in GC patients were much higher compared to normal healthy subjects and that the expression levels of VIP were positively correlated not only to those of VPAC1 expression, but also to metastatic potentials and poor prognosis of GC patients. The serum VIP concentrations are usually increased in the patients with GI cancers (Hogan PG, 2017), but the underlying mechanisms are still obscure. The much higher expression of VIP in GC tissues prompted us to test the hypothesis that GC cells might secrete VIP via an autocrine regulatory mechanism to raise its serum levels. Indeed, we found that VPAC1 activation enhanced VIP expression in GC cells, which was significantly attenuated by Ca^2+^ chelation, indicating a positive regulation of VIP gene transcription by Ca^2+^ signaling in GC cells. Previous studies showed that CREB was involved in Ca^2+^-mediated activation of brain-derived neurotrophic factor gene promoters (Tabuchi A *et al*, 2002). Similarly, VIP gene promoter contains a cAMP-responsive element (CRE) located 70 bps upstream of the transcription initiation site (Hahm SH et al, 1998), which is supported by our observation that the knockdown of CREB attenuated VIP-induced expression of the neuropeptide itself. Therefore, it is most likely that VIP expression is enhanced by Ca^2+^ signaling through promoting the binding of CREB to CRE of VIP gene. Furthermore, VIP-induced Ca^2+^ entry through TRPV4 channels is also likely critical for the exocytosis (Schwarz EC *et al*, 2013), which is consistent with the previous reports that activation of VPAC1 induced a transient membrane depolarization that might trigger Ca^2+^ signaling to promote cytokine secretion (Tompkins JD et al, 2010; Liu F *et al*, 2015). Therefore, the present study elucidates the detailed mechanism underlying the increase of serum VIP in the patients with GI cancer.

In conclusion, we have identified a novel molecular pathological mechanism in which VIP promotes GC progression through the VPAC1/TRPV4/Ca^2+^ signaling cascade, which in turn activates VIP expression and secretion *via* CREB to worsen GC progression (Fig 7L). Increased expressions of VIP/VPAC1 have potentially diagnostic and prognostic significance for gastric cancer, and targeted inhibition of VIP/VPAC1 expression and its downstream signaling may be a novel strategy for gastric cancer diagnosis and therapy.

## Materials and Methods

### Ethics statement

Informed consent was obtained for all patients involved in this study. The use of the clinical specimens was approved by Clinical Research Ethics Committee of Third Military Medical University. All animal experimental procedures were performed according to the “Guide for the Care and Use of Laboratory Animals” published by National Institutes of Health and Ethics Committee of the Third Military Medical University.

### Cell culture

GES-1, MKN74, SGC7901, HGC-27, MKN45 and BGC823 were obtained from the Chinese Academy of Sciences (Shanghai, China). GES-1 is an immortalized normal human gastric epithelial mucosa cell line, while MKN74, SGC7901, HGC-27, MKN45 and BGC823 are human GC cell lines, which are commonly used to study GC. Cells were maintained in Dulbecco’s Modified Eagle’s Medium (DMEM) (Hyclone, Waltham, MA, USA) supplemented with 10% fetal bovine serum at 37 °C in a humidified atmosphere containing 5% CO_2_. The cells were used for experiments after they reached to 70-80% confluence.

### Clinical specimens and Immunohistochemistry

Tissue samples were obtained under sterile conditions from patients with primary GC who underwent surgery at the Department of Surgery, Xingqiao Hospital, Third Military Medical University, Chongqing, China. A diagnosis of GC was confirmed on histological examination. Tissue section was obtained from each patient and blocked with 2.5% hydrogen peroxide in methanol. Then, tissues were incubated with rabbit anti-VPAC1 (Millipore, Temecula, CA, USA), anti-VIP (Santa Cruz, CA, USA), or anti-TRPV4 (Millipore, Temecula, CA, USA) antibodies at 4°C overnight, washed and then followed by incubation with a horseradish peroxidase anti-rabbit IgG antibody. Immunostaining was developed by incubation with diaminobenzidine. The staining results were observed by light microscopy, and ImagePro Plus (Media Cybernetics, MD, USA) was used to quantitatively score the tissue sections. Briefly, images of all tissue cores were acquired at the same time with a constant set of imaging parameters on the microscope and imaging software. The images were then subjected to optical density analysis by ImagePro Plus software. The intensity range selection was based on histograms, with the intensity (I), saturation (S), and maximum hue (H) set at a range in which most of the brown diaminobenzidine color could be quantified. After defining the AOI (area of interest), the mean optical density of the selected area [integrated optical density (IOD)/unit area] was determined by the software and used to represent the immunoreactivity of the candidate protein within the tumor tissue. The acquired score of the optical density was normalized and subjected to ImagePro Plus for analysis.

### Quantitative real-time PCR

Total RNA was extracted by RNAiso plus (Takara, Japan). The RNA was reverse transcribed to cDNA using PrimeScript RT-polymerase (Takara, Japan). Quantitative PCR was performed using primers specific for VPAC1, VIP and TRPV4. The gene GAPDH was used as an internal control. All quantitative PCR assays were conducted using SYBR Premix EX Taq (Takara, Japan) and an ABI detection system (Applied Biosystems, Foster City, CA, USA). All reported results are average ratios of three independent experiments.

### Western blotting

Cells and tissue samples were lysed with a lysis buffer and centrifuged at 12,000 g for 15 min to remove insoluble materials. The protein concentration was measured using a Bradford Protein Assay Kit (Beyotime, Beijing, China). The lysates were separated by SDS-PAGE (10%). Resolved proteins were transferred onto a PVDF membrane (Millipore, Billerica, MA, USA). Membranes were blocked in the blocking buffer, and then incubated with primary antibodies: anti-VPAC1, anti-VIP, anti-TRPV4, anti-p-ERK1/2 (Cell Signaling Technology (CST), Beverly, MA, USA), anti-p-JNK (CST, MA, USA), anti-pSer552 (CST, MA, USA), anti-pSer675 (CST, MA, USA), anti-β-catenin (Abcam, Cambridge, MA, USA) and mouse monoclonal anti-GAPDH (Abcam, Cambridge, MA, USA). These were followed by incubation with appropriate secondary antibodies conjugated with horseradish peroxidase and detection using chemiluminescence (Millipore, Temecula, CA, USA).

### Scratch Assay

Cells from each group were seeded into 24-well plates at a density that reached 80% confluence as a monolayer at 24 h. The monolayer was gently scraped in a straight line with a 10 μl pipette tip to create a “scratch”. The scratch created was of a similar size in the different experimental conditions to minimize possible variations caused by the difference in the initial width. After scratching, the well was gently washed with medium to remove the detached cells. After different treatments as described, images were acquired at 0 h and 24 h under a microscope. In each group, at least 3 parallel wells were utilized for testing.

### Boyden Chamber Assay

The Boyden chamber assay was performed to evaluate the ability of cells to migrate, which makes use of a chamber composed of two medium filled compartments separated by a micro porous membrane (BD Biosciences, San Diego, CA, USA). The lower well contained medium with 5% (*vol*/*vol*) serum. Cells were placed in the upper chamber and treated with VIP or inhibitors, such as RN1734 or other treatments as described. Cells were allowed to migrate through the pores of the membrane into the lower compartment. After 24 h, cells that had migrated onto the lower surface were stained with crystal violet. The number of cells that had migrated to the lower side of the membrane was evaluated under a microscope (Olympus, Japan). Triplicate experiments with triplicate samples were performed.

### Cell invasion assay

Briefly, 130 μl of Matrigel Basement Membrane Matrix (BD Biosciences, San Diego, CA, USA) was added to each well of the precooled 24-well transwell chamber (BD Biosciences, San Diego, CA, USA). Pipette tips and Matrigel solution were kept cold throughout to avoid solidification. The plate was incubated at 37 °C for 1 h to allow the matrix solution to solidify. A total of 1 × 10^4^ cells in a final volume of 500 μL of basal medium (DMEM) were seeded onto the surface of each well containing the polymerized matrix. Cells were pretreated with desired inhibitor or with vehicle alone and stimulated with the specific agonist VIP for 24 h at 37 °C. Cells remaining on the upper surface of the filters were removed by wiping with cotton swabs, and invading cells were fixed with 4% paraformaldehyde, stained with crystal violet and dried. The invasive cells were evaluated under a microscope (Olympus, Japan). Triplicate experiments with triplicate samples were performed.

### Measurement of [Ca^2+^]_cyt_ by calcium imaging

GC cells cultured on coverslips were loaded with 5 μM Fura-2AM (Invitrogen, NY, USA) in a physiological salt solution (PSS) at 22°C for 50 min and then washed in PSS for 30 min. Then, the coverslip was placed in a standard perfusion chamber on the stage of an inverted fluorescence microscope (Nikon, Japan). Images of Fura-2 fluorescence with excitation at 340 and 380 nm were captured alternately over time using an intensified CCD camera (ICCD200) controlled by the MetaFluor software (Universal Imaging Corporation, Downingtown, PA). F340/380 ratios in areas of interest were used to represent [Ca^2+^]_cyt_. PSS contained (in mM): 140 Na^+^, 5 K^+^, 2 Ca^2+^, 149 Cl^-^, 10 HEPES, and 10 glucose, pH 7.4. For the Ca^2+^-free solution, Ca^2+^ was omitted from and 0.5 mM EGTA was added to PSS to prevent possible Ca^2+^ contamination. All inhibitors or treatments were applied as described.

### Electrophysiological assessment of TRPV4 currents

Whole-cell voltage clamp recordings were made from cultured GES-1 and MKN-45 cells seeded on glass coverslips (~40% confluence) using an EPC10 amplifier (HEKA Elektronik). Patch electrodes with resistances of 3-5 MΩ were pulled from borosilicate micropipettes (Sutter Instrument) and filled with a pipette solution containing (in mM): 130 K-methanesulfonate, 7 KCl, 0.05 EGTA, 10 HEPES, 1 Na_2_-ATP, 3 Mg-ATP, 0.05 Na_2_-GTP, pH 7.3 adjusted with KOH, 300 mOsM. The extracellular solution (ECS) contained (in mM): 140 NaCl, 5 KCl, 2 CaCl_2_, 1 MgCl_2_, 10 HEPES, 10 Glucose, pH 7.4 adjusted with NaOH, 310 mOsM. Cells were held at 0 mV. Voltage ramps from −100 to 100 mV in 500 ms were applied repetitively every 2 sec while the recorded cell was continuously perfused with ECS. VIP and RN1734 were applied at desired time periods through perfusion. All recordings were conducted at room temperature (~23°C).

### Immunofluorescence staining

GC cells were grown on coverslips, fixed with 4% paraformaldehyde (w/v) for 30 min, and then permeabilized with 0.5% Triton X-100 in phosphate-buffered saline (PBS) and blocked with bovine serum albumin. The cells were incubated with primary rabbit anti-VIP or anti-TRPV4 (Millipore, Temecula, CA, USA) at 4 °C overnight. After careful washing with PBS, the cells were incubated with secondary antibodies conjugated to Cy3. Fluorescent images were observed and analyzed with a laser scanning confocal microscope (Olympus, Japan).

### ELISA assay

VIP levels were determined by sandwich ELISA using human VIP ELISA Kit (Biocompare, South San Francisco, CA, USA) according to the manufacturer’s instructions. Briefly, cells were treated as indicated, and 100 μl of the culture medium or 40 μl of serum were transferred into microplate wells precoated with a monoclonal antibody specific for VIP, and incubated for 2 h at 37 °C. An enzyme-linked polyclonal antibody was then added to the wells and incubated for 1 h at 37 °C to sandwich the VIP ligand. After careful wash, a substrate solution was added and the intensity of the color was determined using an automicroplate reader (Thermo Scientific, Carlsbad, CA, USA). Each experiment was conducted twice and each sample point was assessed in triplicate.

### In vivo metastasis assay

MKN45 cells, untransfected or transfected with the lentiviral vectors Lv-sh-NC, Lv-sh-VPAC1, Lv-sh-VIP, or Lv-sh-TRPV4 were used. For the *in vivo* lung metastasis assay, 1×10^6^ transfected cells were suspended in 200 μl PBS for each mouse. The cells were injected into nude mice (five mice per group, five-week-old male BALB/c-nu/nu mice) through the tail vein. Then VIP or PBS was injected through tail vein once a day. After 12 weeks, the mice were sacrificed. Each lung was divided into five blocks, embed in paraffin, and the paraffin mass was cut into five sections, which were stained with hematoxylin and eosin. The metastatic nodules were observed under a microscope and calculated. For the peritoneal dissemination assay, MKN45 cells were transfected as indicated, and 1×10^6^ transfected cells were injected into the abdominal cavity of nude mice (five mice per group, five-week-old male BALB/c-nu/nu mice). Then, VIP or PBS was injected into the abdominal cavity once a day. Six weeks later, the mice were killed, and the nodules were observed and counted.

### Statistical Analysis

Data are expressed as means ± SEM. A two-tailed Student’s t-test or Mann-Whitney test was used to determine the differences between groups. The Kaplan-Meier curves were plotted to assess the effect of VIP/VPAC1 expression on survival, and survival curves were compared using the log-rank test. The χ^2^ test was used to test possible associations between VIP/VPAC1 expression and clinicopathological factors. The correlation coefficients were determined using Pearson’s rank correlation test. Multivariable Cox proportional hazards regression models were used to assess the prognostic significance of VIP/VPAC1 expression. IBM SPSS Statistics V.22 software (IBM, Armonk, New York, USA) was used for statistical analysis, and the graphs were generated using GraphPad Prism 6.0 (Graphpad Software Inc, CA, USA).

## Acknowledgments

This work was supported by the National Key Research and Development Program of China (No. 2016YFC1302200 to HD), National Natural Science Foundation of China (No. 81602577 to BT), and Basic Science and Frontier Technology Research Project of Chongqing (No. cstc2017jcyjAX0149 to BT).

## Competing financial interests

The authors declare no competing financial interests.

## Author Contributions

HD and SY conceived of the study; BT and MXZ designed the experiments; BT, JW, XS, JL, RX, TD and YX performed the experiments; BT and JW performed analysis and interpretation of data; HD and BT wrote the manuscript; MXZ, JMC and SY critically reviewed the manuscript.

## The Paper Explained

### Problem

Gastric cancer is the second leading cause of cancer-related death worldwide. Although the diagnosis and treatment have been improved over the past decades, the prognosis remains poor due to its early invasion and metastasis. Tumor invasion and metastasis is a complex biological process, and growing evidence show that the interaction between cancer cells and the surrounding microenvironment is critical for tumor metastasis. However, the molecular mechanisms of the invasion and metastasis of gastric cancer are still poorly understood.

### Results

We revealed that VPAC1 was overexpressed in GC tissues and was significantly associated with the depth of invasion, advanced TNM stage and distant metastasis. High expression of VPAC1 in GC was an independent prognostic predictor and the GC patients with high VPAC1 expression usually had poor survival. Mechanistically, VPAC1 activation by VIP markedly induced TRPV4-mediated Ca^2+^ entry and GC progression. VPAC1/TRPV4/Ca^2+^ signaling in turn enhanced the expression and secretion of VIP in GC cells, enforcing a positive feedback regulation in GC progression. Our findings suggest that VPAC1 has potentially diagnostic and prognostic significance for GC, and targeting VPAC1 and its downstream signaling could attenuate GC metastasis.

### Impact

Our results highlight the importance of VPAC1/TRPV4/Ca^2+^ signaling in gastric cancer development, and provide a novel strategy for developing new drugs targeting this pathway for gastric cancer treatment.

